# Tuning the H_2_ Production Activity of ToHydA by Molecular Simulation-Informed Protein Engineering

**DOI:** 10.64898/2025.12.18.695218

**Authors:** Chandan K. Das, Subhasri Ghosh, Shuvankar Naskar, Thomas Happe, Lars V. Schäfer

## Abstract

In the context of bio-hydrogen production, the O_2_-stable Group B [FeFe]-hydrogenase from *Thermosediminibacter oceani* has attracted significant interest due to its distinctive “TSCCCP” motif near the active site, which contains an additional (third) cysteine residue that is absent in the “TSCCP” motif of standard Group A hydrogenases. The precise role of the additional cysteine residue in H_2_ production has remained an open question. In this study, we sought to contribute to the understanding of this cysteine’s role in H_2_ production by combining molecular dynamics (MD) simulations, site-directed mutagenesis, biochemical assays, and Fourier-transform infrared (FTIR) spectroscopy. Remarkably, a cysteine-to-serine exchange variant (TSSCCP) demonstrated enhanced H_2_ production activity without compromising the O_2_-stability of ToHydA, offering new insights into its functional dynamics.

---

The efficient bio-production of molecular hydrogen by [FeFe]-hydrogenase, a redox metalloenzyme, has attracted significant attention in the context of the growing energy crisis.^1^ The enzyme’s active site, known as the H-cluster, composed of a unique di-iron ([2Fe]_H_) cluster and a [4Fe-4S] ([4Fe]_H_) cubane cluster connected through a cysteine residue, catalyzes the hydrogen production from protons diffusing into the active site, presumably through proton transporting residues.^2–4^ These protons are transferred to the nitrogen atom of the azadithiolate (N_adt_) bridge of the [2Fe]_H_ cluster through a key cysteine residue, often referred to as the proton-transporting cysteine (here named as reactive cysteine due to its key role in catalysis).^4–7^ By accepting two electrons, an H_2_ molecule is generated at the distal iron site (Fe_d_) of the [2Fe]_H_ cluster.^8,9^[FeFe]-hydrogenases are usually highly susceptible to oxidative degradation under oxygen-rich conditions, limiting their stability and practical application.^11–17^

In our previous study, we characterized ToHydA, a Group B [FeFe]-hydrogenase from *Thermosediminibacter oceani*, which exhibits remarkable stability even after prolonged exposure to oxygen, highlighting the potential of ToHydA as a robust biocatalyst for hydrogen production.^10^ ToHydA features a TSCCCP motif near the H-cluster, where one cysteine (C213) coordinates the [4Fe]_H_ cluster, while the reactive cysteine (C212) facilitates proton transfer to the adt bridge (Figure 1). The remaining cysteine (referred to as additional cysteine, C211) is a distinctive characteristic of Group B (M2a) hydrogenases.^18^ The deletion of this cysteine (dC211 variant) has been demonstrated to enable the continued functionality of To-HydA. Introducing a proline residue four positions upstream to compensate for the missing C211 (mimicking the protein sequences of native group A hydrogenases) restores wild-type activity. However, the role of this cysteine in the catalytic cycle is unclear. Here, a combined approach involving molecular dynamics (MD) simulations, site-directed mutagenesis, biochemical assays and ATR-FTIR spectroscopy was employed to investigate the role of C211 in ToHydA’s catalytic activity.

**Figure 1.**
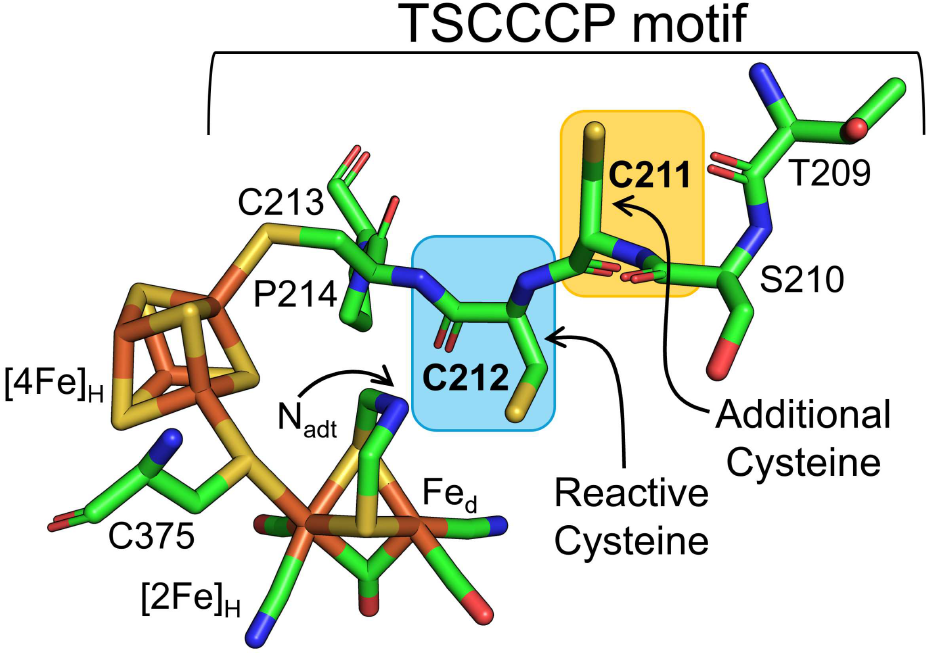
Structural view of the active site of ToHydA (generated using AlphaFold).^10^ The figure highlights the TSCCCP motif, with the reactive cysteine and the additional cysteine marked by blue and yellow squares, respectively.

First, we performed microsecond timescale all-atom MD simulations of wild-type (WT) ToHydA. For analysis, we used the final 500 ns of each of the three replicate simulations, yielding a total of 1500 ns of MD simulation time for each system (see Methods). In the MD simulations of the WT enzyme, we observed a low occurrence of the hydrogen bond between N_adt_ and C212:H_γ_ (Figure 2A, B), which is an essential interaction for efficient proton transfer. This broken H-bond suggests reduced hydrogen production activity. In addition, the simulations also revealed that the C211 side chain occasionally forms an H-bond with the backbone carbonyl group of S234 (Figure 2C). These two interactions are correlated, in the sense that the formation of the C211:H_γ_…S234:O hydrogen bond appears to facilitate the formation of the C212:H_γ_…N_adt_ hydrogen bond due to an anti-conformation of the side chains of C211 and C212 (Figure 2). Taken together, we thus hypothesize that the C211:H_γ_…S234:O hydrogen bond promotes the formation of the C212:H_γ_…N_adt_ hydrogen bond. To test that hypothesis, we performed MD simulations of the C211S variant, in which the cysteine residue at position 211 is replaced by a serine, which is more polar than cysteine and might thus be a stronger H-bond donor. Indeed, the simulations revealed a significant increase in the formation of the S211:H_γ_…S234:O hydrogen bond, which in turn enhanced the occurrence of the C212:H_γ_…N_adt_ hydrogen bond that might promote hydrogen production (Figure 2B and C, magenta dots).

**Figure 2.**
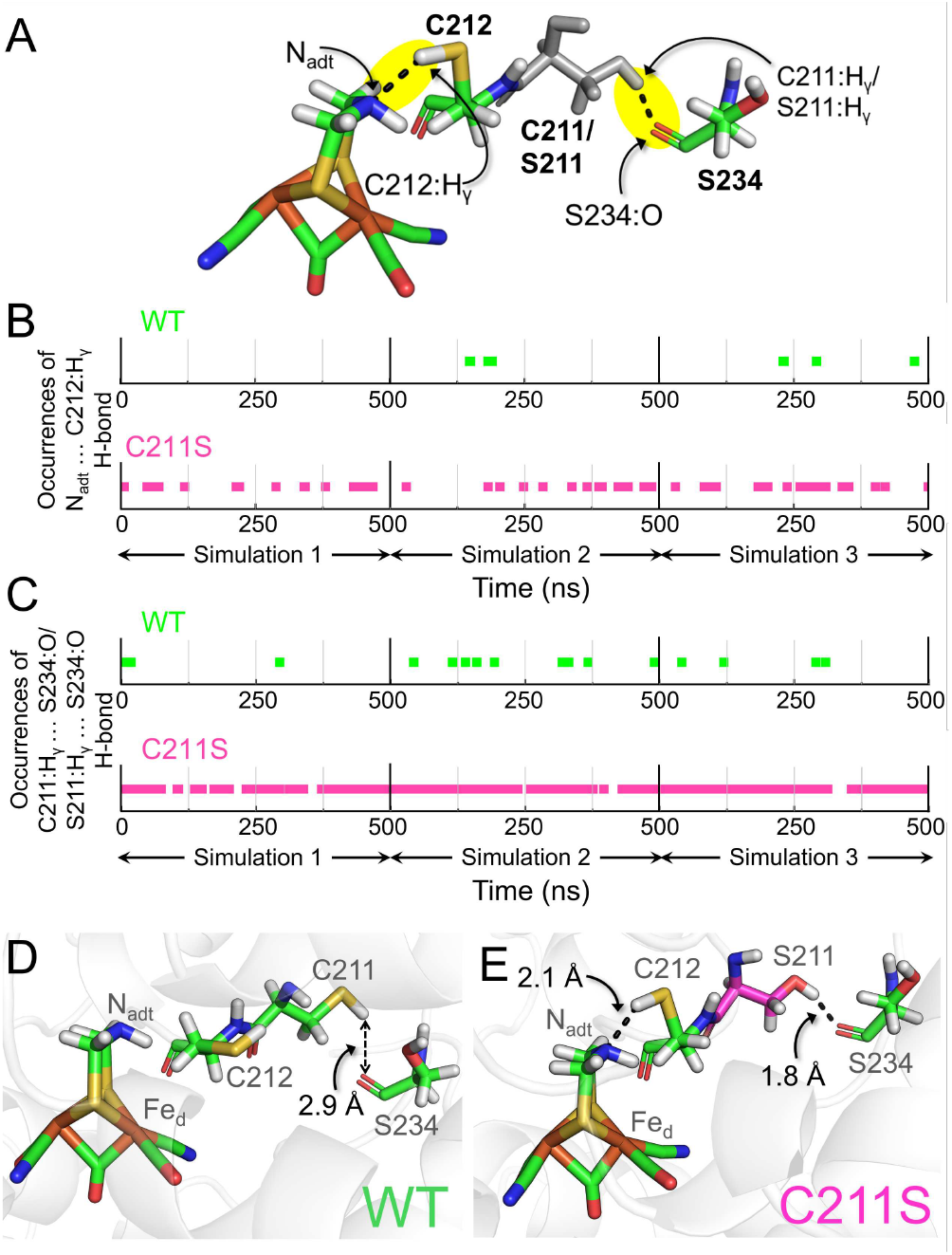
A) Schematic representation of active site highlighting C212:H_γ_…N_adt_ and C211:H_γ_/S211:H_γ_…S234:O hydrogen bonds. B) and C) The occurrences of these hydrogen bonds in three independent 500ns simulations of ToHydA WT and C211S variant. D) and E) Representative MD snapshots of ToHydA WT and the C211S variant.

On the basis of our MD results, we decided to experimentally investigate the ToHydA WT and the C211S variant. The proteins were expressed and purified following previously described protocols (SI experimental details).^10,19^ The WT displays an *in-vitro* H_2_ production activity of 100 µmol H_2_.min^-1^.mg^-1^, while the C211S variant demonstrated a significantly higher activity of 170 µmol H_2_.min^-1^.mg^-1^ (Figure 3A). To investigate the catalytic states of both proteins, ATR-FTIR spectroscopy was employed. The as-isolated WT and C211S proteins show very similar peaks and primarily displayed the H_inact_ and H_trans_ states, along with minor contributions from reduced states (Figure 3B and 3C, upper panels). When exposed to H_2_, the WT showed an increased peak at 1884 cm^-1^, which is linked to a terminal CO ligand and is usually seen with the H_red_ or H_sred_ state. While distinguishing between these two states was beyond the scope of this study, we collectively refer to all peaks observed between 1955 cm^-1^ and 1884 cm^-1^ (except H_ox_ peaks) as “reduced states”. The 1884 cm^-1^ peak, being the most prominent, serves as a marker for the accumulation of reduced states. In the C211S variant, H_2_ treatment led to a notable increase in the intensity of the 1884 cm^-1^ peak, along with additional peaks at 1909, 1922, and 1955 cm^-1^. This observation suggests a higher accumulation of reduced states in the C211S variant compared to the WT. It is worth noting that the reactive cysteine (C212) can adopt two different conformations, which directly influence the accumulation of certain catalytic states. In the catalytically relevant conformation of C212, the C212 side chain is oriented away from the Fe_d_ atom and forms a hydrogen bond with the N_adt_ bridgehead, thereby facilitating efficient proton transport to the active site (Figure 3D). Conversely, in the other conformation the C212 sidechain is located closer to the Fe_d_ (Figure 3E). The latter conformation eventually leads to the formation of the H_inact_ and H_trans_ states, while the former is adopted in catalytic intermediates such as the H_ox_ state and the reduced states.^20,21^ In the C211S variant, the higher accumulation of reduced states indicates an increased presence of the first conformation, leading to increased catalytic activity. These interpretations are in line with the predictions of the MD simulations. The C211S substitution did not compromise the O_2_ stability of ToHydA. Following 15 minutes of O_2_ exposure, the FTIR spectra of the C211S variant revealed the accumulation of the H_inact_ state without any detectable protein degradation (SI Figure S1).

**Figure 3:**
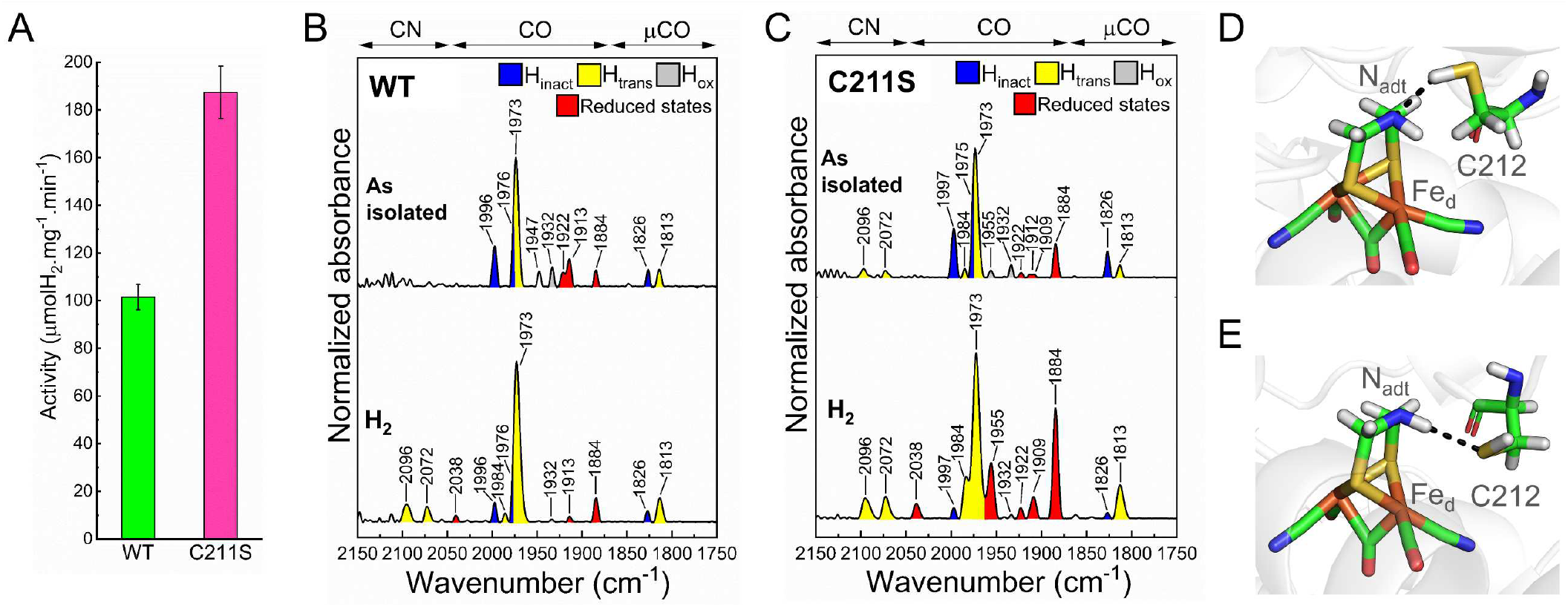
A) *In-vitro* H_2_ production activity of (anaerobic) ToHydA WT and C211S variant, showing higher activity for C211S variant. Technical and biological replicates are shown in SI Figure S2. B) and C) FTIR spectra of ToHydA WT and C211S variant in the as-isolated state (upper panel) and after H_2_ purging (25 L/min) (lower panel). Under H_2_ purging condition, C211S variant accumulates reduced states compared to ToHydA WT. All gas purging experiments started from as-isolated state. The spectra of the proteins are normalized to the second amide band (1535 - 1545 cm^−1^). 0.4 - 0.5 mM protein sample was prepared in 100 mM Tris/HCl buffer at pH 8.0 containing 2mM NaDT. D) and E) Two distinct hydrogen-bonded conformations of C212.

As a negative control of our mechanistic hypothesis, we substituted the additional cysteine with alanine in the ToHydA C211A variant. This variant clearly lacks the hydrogen bonding interaction with S234 (Figure 4). The C212:H_γ_…N_adt_ hydrogen bond is almost entirely absent, highlighting how the interaction between the side chain of residue 211 and the backbone carbonyl group of S234 directs the formation of the C212:H_γ_…N_adt_ hydrogen bond (Figures 3A and 3B). Indeed, the C211A variant exhibited 50% of the WT’s in vitro catalytic activity (SI Figure S2). Additionally, the reduced state peaks in the H_2_-treated FTIR spectra were diminished compared to the WT (Figure 4C). The reduced catalytic activity, along with the lower accumulation of reduced state peaks in the FTIR spectra of C211A, supports the proposed hypothesis.

**Figure 4.**
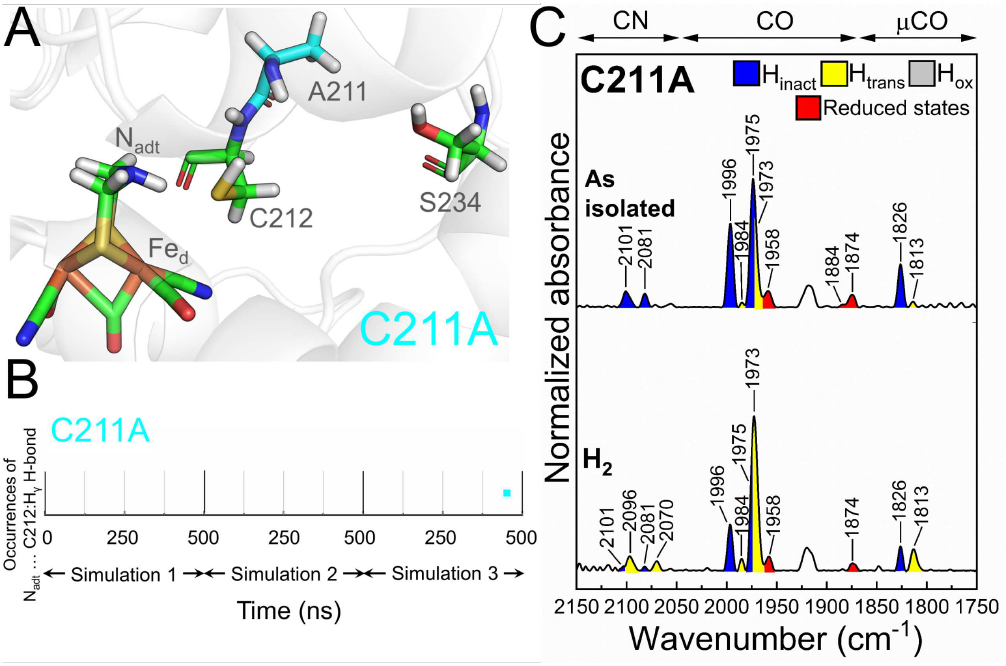
A) MD snapshot of ToHydA C211A variant. B) The occurrences of hydrogen bond between N_adt_ and C212:H_γ_ in three independent 500 ns simulations of ToHydA C211A variant. C) FTIR spectra of ToHydA C211A in the as-isolated state (upper panel) and after H_2_ purging (25 L/min) (lower panel). All gas purging experiments started from as-isolated state. The spectra of the proteins are normalized to the second amide band (1535 - 1545 cm^−1^). 0.4 - 0.5 mM protein sample was prepared in 100 mM Tris/HCl buffer at pH 8.0 containing 2mM NaDT.

To conclude, in this study, we investigated the function of the additional cysteine near the H-cluster in Group B hydrogenases. Previously, Fasano et al. proposed that the side chain of this additional cysteine could replace one of the two conserved water molecules in the proton transport pathway (identified in Group A hydrogenases)^4^ of CpIII, another Group B [FeFe]-hydrogenase.^22^ According to their hypothesis, the additional cysteine side chain might assume the role of maintaining the hydrogen-bonding network with the reactive cysteine and the remaining water molecule. However, we do not observe an orientation of C211 that is optimal for proton shuttling. Instead, C211 influences the orientation of the C212 side chain, which aligns with the findings from the C211S and C211A variants. Furthermore, the C211S variant could open new avenues for enhancing H_2_ production activity in O_2_-stable [FeFe]-hydrogenases through protein engineering.

## ASSOCIATED CONTENT

### Supporting Information

The supporting information includes experimental and MD simulation details, FTIR spectra of O_2_-treated C211S, technical and biological replicates of biochemical assays, SDS-PAGE of the isolated proteins and primer sequence for generation of site-directed mutagenesis variants.

## AUTHOR INFORMATION

### Author Contributions

The manuscript was written through contributions of all authors. All authors have given approval to the final version of the manuscript. All authors performed research and/or analyzed data. C.K.D. conducted all the MD simulations. S.G. expressed and isolated the proteins and performed the biochemical experiments. S.G. and S.N. performed ATR-FTIR spectroscopy. C.K.D., S.G., S.N., L.V.S., and T.H. wrote and/or edited the manuscript. ‡These authors contributed equally.

### Notes

The authors declare no competing financial interest.

## ACKNOWLEDGMENT

S.G. thanks the Deutscher Akademischer Austauschdienst (DAAD) for funding her doctoral scholarship. This project received funding from the Deutsche Forschungsgemeinschaft (DFG) under Germany’s Excellence Strategy – EXC 2033 – 390677874 – RESOLV. T.H. thanks the Volkswagen Stiftung (Az 98621). We thank Shanika Yadav and Ulf-Peter Apfel from the Faculty of Chemistry and Biochemistry at Ruhr University Bochum for their synthesis and provision of the [2Fe]^MIM^ used in *in-vitro* maturation.

## Notes

### Competing Interest Statement

The authors have declared no competing interest.

