## Supporting Information for "Tuning the H_2_ Production Activity of ToHydA by Molecular Simulation-Informed Protein Engineering"

#### Molecular dynamics simulation details

All molecular dynamics (MD) simulations were performed using the GROMACS software package (version 2021.1).[1] The initial structure of the ToHydA WT [FeFe]-hydrogenase was obtained from previous work.[2] In the C211S and C211A variants, the C211 residue was substituted with serine and alanine, respectively. The same MD protocol as in previous work was applied for these simulations.

The simulations were carried out using the CHARMM36 protein force field in combination with the CHARMM-specific TIP3P water model.[3] The study specifically examined the $H_{ox}$ state of ToHydA [FeFe]-hydrogenases. Force field parameters for the H-cluster and accessory FeS clusters, along with the coordinating cysteine and histidine residues, were derived from the work of Chang et al.,[4] with further adjustments based on recommendations by McCullagh and Voth.[5]

Each protein was fully solvated in a system containing 16,519 water molecules, and the net charge was neutralized by adding 17 $Na^+$ ions. The simulations were conducted using periodic boundary conditions within dodecahedron-shaped simulation boxes. The total number of atoms in the system was 56,208 for the C211S variant and 56,207 for the C211A variant.

Prior to conducting MD simulations, the systems underwent energy minimization using 10,000 steps of steepest descent. This was followed by stepwise equilibration with harmonic position restraints applied to specific atom groups. The equilibration process started with an NVT simulation, during which the system's temperature was gradually increased from 0 K to 300 K over 0.2 ns. During this phase, position restraints with force constants of 1,000 kJ/mol/nm$^2$ were applied to all non-hydrogen atoms of the protein and FeS clusters. Equilibration then continued for 2.5 ns in the NPT ensemble. For the first 0.5 ns of this phase, the same position restraints were maintained on all non-hydrogen atoms of the protein and FeS clusters. During the remaining 2.0 ns of NPT equilibration, restraints were limited to the protein backbone atoms, allowing the side chains, FeS clusters, and surrounding water molecules to adjust and relax. Finally, three production simulations of 1,000 ns each were performed in the NPT ensemble at 300 K for each of the three protein systems. These simulations began with the equilibrated systems, using different random seeds to assign initial atomic velocities based on a Maxwell-Boltzmann distribution. For analysis, the last 500 ns of each simulation were used.

During the MD simulations, the temperature was kept constant at 300 K using the velocity rescaling thermostat developed by Bussi and colleagues, with a coupling time constant of 0.1 ps.[6] The pressure was maintained at 1 bar through the use of an isotropic weak coupling Berendsen barostat,[7] which had a coupling time constant of 2 ps and a compressibility of $4.5 \times 10^{-5}$ bar$^{-1}$. Short-range Coulomb and Lennard-Jones 6,12 interactions were calculated using a buffered Verlet pair list.[8] Potentials were smoothly tapered to zero at a cutoff distance of 1.2 nm, with forces gradually transitioning to zero between 1.0 and 1.2 nm. Long-range electrostatic interactions were handled using the particle mesh Ewald (PME) method with a grid spacing of 0.12 nm.[9] The LINCS algorithm was employed to constrain all bonds involving hydrogen atoms in the protein,[10] while the SETTLE algorithm was used to fix all internal degrees of freedom for water molecules.[11] This enabled the integration of the equations of motion with a time step of 2 fs. For analyzing hydrogen bonds, a heavy atom distance cutoff of 0.3 nm and an angle cutoff of 30° between the donor and acceptor heavy atoms were applied.

### Experimental details

#### Site-directed mutagenesis

Hydrogenase genes with varied codons were synthesized using QuikChange PCR, based on a previously established protocol.[12] Expression plasmids were generated by amplifying with mismatch primers (refer to SI Table S1). The PCR products were then digested with DpnI endonuclease and introduced into *E. coli* DH5α via the heat shock transformation method. All DNA constructs were verified by sequencing.

#### Expression and purification

The *E. coli* strain BL21 (DE3) ΔiscR3[13] was transformed with a pET21b plasmid containing a codon-optimized ToHydA gene to produce the apo form of ToHydA, which does not include the $[2Fe]_H$ subcluster.[14] Expression and purification were performed under strictly anaerobic conditions without the involvement of the hydrogenase maturases HydE, HydF, and HydG, following established protocols.[2,14] Affinity chromatography using Strep-Tactin high-capacity resin (IBA GmBH) was employed to purify the protein. The Bradford assay was used to determine protein concentration,[15] and SDS-PAGE analysis (see SI Figure S3) verified purity.[16] The purified protein was stored at −80 °C in 100 mM Tris-HCl buffer (pH 8) with 2 mM sodium dithionite (NaDT).

#### *In-vitro* maturation

The active protein was prepared by incubating apoproteins on ice for 1 hour with a 10-fold molar excess of the synthetic $[2Fe]_H$ cofactor mimic ($[2Fe]^{MIM}$ = $[Fe_2[\mu\text{-}(SCH_2)_2NH]\text{-}(CN)_2(CO)_4]^{2-}$)[17] in 100 mM potassium phosphate buffer (pH 6.8) supplemented with 2 mM NaDT, following a previously reported procedure.[18] After the incubation, holoproteins were separated from the unbound $[2Fe]^{MIM}$ mimic using size exclusion chromatography with a NAP 5 column (GE Healthcare). The mature holo-proteins were then stored at −80 °C in 100 mM Tris-HCl buffer (pH 8) containing 2 mM NaDT.

#### $H_2$ production assay

To measure *in vitro* hydrogen production activity, a reaction mixture was prepared in an 8 mL airtight glass vessel. The mixture included 10 mM methyl viologen (MV) as an electron mediator, 100 mM NaDT as a sacrificial electron donor, and 800–1600 ng of holoenzyme in 100 mM potassium phosphate buffer (pH 6.8). The mixture was degassed with 100% argon for 5 minutes and then incubated at 37 °C for 30 minutes. Hydrogen production in the headspace was quantified using gas chromatography (Shimadzu).[19]

ATR−FTIR spectroscopy

Attenuated total reflectance Fourier transform infrared (ATR-FTIR) spectroscopy was performed using a Bruker Tensor II spectrometer (Bruker Optik, Germany) equipped with a 9-reflection ZnSe/Si crystal (Microm ATR Vision, Czitek). All measurements were carried out under anaerobic conditions (2% $H_2$ and 98% $N_2$) at 25°C. The FTIR spectra were recorded in the 4000-1000 $cm^{-1}$ range with a resolution of 2 $cm^{-1}$. For sample preparation, 4 µL of protein sample at a concentration of 0.5 mM stored in 100 mM Tris-HCl buffer (pH 8) containing 2mM NaDT was applied to the ATR crystal.

The "as-isolated" ToHydA spectrum was obtained by drying the sample in a tented environment without the use of an external gas flow. For gas-flushing experiments, the sample was partially dried until the distinctive H-cluster bands became faintly detectable, indicating a spectrum resembling that of the "as-isolated" state, though with reduced intensity and preserved hydration. To maintain hydration and prevent further drying, the protein film was exposed to a humidified gas stream containing $H_2$ (25 L/min for approximately 10 minutes) and $O_2$ (air, 2 L/min for around 15 minutes). Compared to our previous work,[2] a fivefold increase in the $H_2$ flow rate (previously 5 L/min flowrate of $H_2$ used) was employed to enhance protein reduction without the use of methyl viologen. This adjustment helps mitigate the formation of the $H_{inact}$ state and promotes higher accumulation of reduced states.

### Supporting figures and tables

#### Figure S1. ATR-FTIR spectra of $O_2$-treated ToHydA WT and C211S

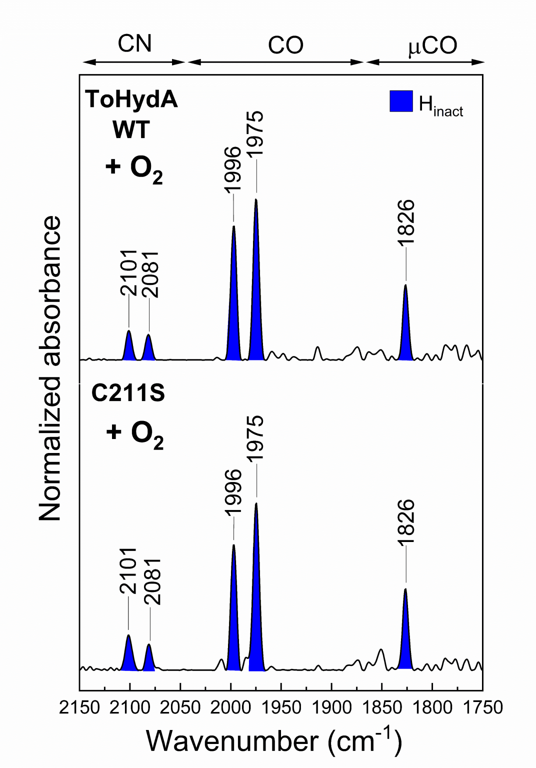

**Figure S1:** FTIR spectra of $O_2$-treated (air, 2 L/min) ToHydA WT (upper panel) and the C211S variant (lower panel) reveal no difference in $H_{inact}$ accumulation for the C211S variant. All gas purging experiments started from as-isolated state. The spectra of the proteins are normalized to the second amide band (1535–1545 $cm^{-1}$). 0.4–0.5 mM protein sample was prepared in 100 mM Tris-HCl buffer (pH 8) containing 2 mM NaDT.

Figure S2. *In-vitro* $H_2$ production activity of ToHydA WT and variants

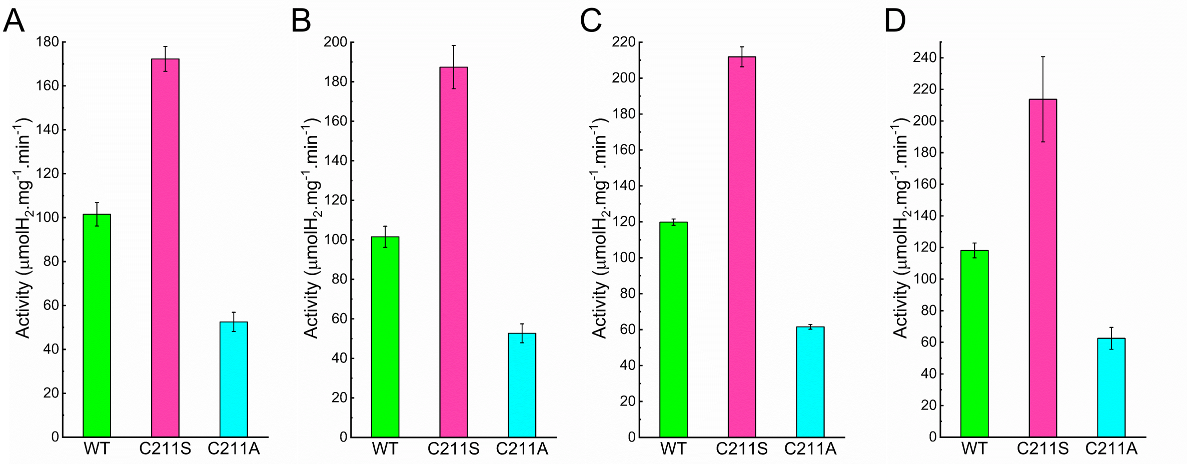

**Figure S2:** In vitro MV-mediated anaerobic $H_2$ production activity of ToHydA WT and its variants was assessed. Panels A and B represent technical replicates derived from a single protein preparation (Biological replicate 1), measured on different days. Similarly, panels C and D show technical replicates from an independent protein preparation (Biological replicate 2), also measured on separate days. Each dataset represents the mean of three measurements, with error bars indicating the standard deviation.

Figure S3. SDS PAGE of ToHydA variants after purification

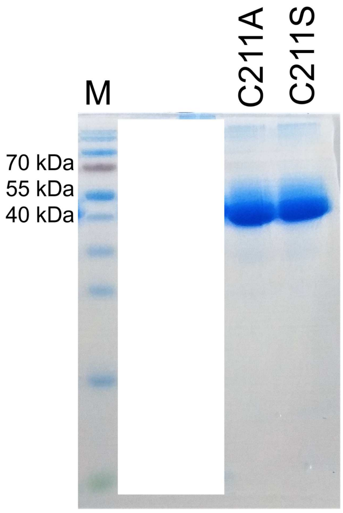

**Figure S3:** The left lane shows protein marker. The middle and right lane represent ToHydA variants C211A and C211S, respectively.

Table S1. Generation of site-directed mutagenesis variants of ToHydA. Primers used for QuikChange PCR to generate variants of ToHydA.

| | |
|---|---|
| **C211S_for:** | CCAGCAGCTGTTGTCCGCCTTTTGTTC |
| **C211S_rev:** | GGACAACAGCTGCTGGTAATCATGAATTTATC |
| **C211A_for:** | CCAGCGCGTGTTGTCCGCCTTTTGTTC |
| **C211A_rev:** | GGACAACACGCGCTGGTAATCATGAATTTATC |
